# Differential modulation of crassulacean acid metabolism according to macronutrient deficiencies in C_4_ *Portulaca oleracea*

**DOI:** 10.64898/2026.07.24.740630

**Authors:** Renata Callegari Ferrari

## Abstract

- C_4_ photosynthesis and the crassulacean acid metabolism (CAM) rarely co-evolved in a single lineage, but *Portulaca* can switch from C_4_ to CAM under drought stress. Little is known about CAM responses to nutrient availability, hence the goal of this work was to assess the influence of macronutrients over C_4_-CAM.
- *P. oleracea* was grown hydroponically and subjected to treatments (+/− PEG) for: nitrate deficiency (−NO_3_^−^), ammonium (NH_4_^+^), NO_3_^−^ + NH_4_^+^, magnesium (−Mg), phosphorus (−P), calcium (−Ca), potassium (−K), and sulphur (−S) deficiencies, and salt stress. Samples were monitored for diurnal titratable acidity (ΔH^+^), osmotic potential, and gene relative expression for core C_4_/CAM and signaling genes.
- -NO_3_^−^ induced CAM even without PEG, a process probably without the mediation of abscisic acid (ABA). Notably, −P showed a trend to induce CAM without PEG and −Ca prevented CAM induction even with PEG. Salt stress induced CAM, and NH_4_^+^ was not toxic for *P. oleracea*. Other treatments showed less conspicuous responses.
- This work brings an unprecedented overview of the nutrition of C_4_ and CAM, suggesting perspectives for deepening the study of C_4_-CAM. Understanding the molecular mechanisms and environmental signaling of C_4_-CAM is essential for realizing the evolution of two CCMs in a single plant.

## Introduction

Carbon concentrating mechanisms (CCMs) evolved ca. 30 million years ago to increase availability of CO_2_ close to Rubisco and reduce photorespiratory rates (Edwards & Ogburn, 2012). Photorespiration usually increases when O_2_ competes with CO_2_ for the same binding site at Rubisco, and rates can significantly increase when plants are subjected to abiotic stresses, reducing photosynthetic yields (Walker *et al*., 2024). Two CCMs are the most widespread across land plants: C_4_ photosynthesis and the crassulacean acid metabolism (CAM), occurring in ca. 9% of all angiosperm species (Sage *et al*., 2017; Gilman *et al*. 2023). Both rely on a similar biochemical pathway to ensure Rubisco has access to high concentrations of CO_2_ but show different regulation and physiology (Sage, 2002). They operate via a first carboxylation step using phosphoenolpyruvate carboxylase (PPC), which assimilates CO_2_ into a four carbon (C) compound that is then decarboxylated close to Rubisco.

Evolving either C_4_ or CAM involved the recruitment and re-functionalization via gene duplication of enzymes already operating in C_3_ plants, often with non-photosynthetic roles (e.g., PPC) (Christin *et al*., 2014, 2015). Although evolving C_4_ and CAM in the same plant could seem redundant and result in futile cycling with unnecessary production and consumption of metabolic intermediates, more cases of C_4_ plants able to switch to performing CAM during drought stress continue to be reported (Koch & Kennedy, 1980, Ku *et al*., 1981, Zhang *et al*., 2014, Ho *et al*., 2019, Siadjeu *et al*., 2024). Among those, *Portulaca oleracea* is the currently best studied model for the facultative C_4_-CAM photosynthetic switch (Ferrari & Freschi 2019; Ferrari & Reyna-Llorens 2026).

The photosynthetic pathway operating in C_4_ *P. oleracea* under well-watered conditions takes place during the day and in two different cells (Fig. 1) (Ferrari & Reyna-Llorens, 2026). A C_4_-specific PPC gene (*PPC-1E1a’*) combines HCO_3_^−^ from CO_2_ dissociation by β-carbonic anhydrase (βCA) with phosphoenolpyruvate (PEP) in mesophyll cells forming oxalacetate (Fig. 1) (Kanai & Edwards, 1999). oxalacetate is converted to aspartate via aspartate aminotransferase (AspAT) and shuttled to the bundle sheath cells, where it will form malate via NAD-malate dehydrogenase (NAD-MDH) and subsequently will be decarboxylated by NAD-malic enzyme (NAD-ME), releasing CO_2_ close to Rubisco and defining the NAD-ME subtype of C_4_ (Fig. 1) (Alvarez and Maurino 2023). There might also be some contribution of phosphoenolpyruvate carboxykinase (PEPCK) as a second decarboxylase as well (Fig. 1, Lara *et al*., 2003, Alvarez & Maurino, 2023). The remaining pyruvate from the previous reaction is interconverted to alanine in bundle sheath cells by alanine aminotransferase (AlaAT), transported back to mesophyll cells, reconverted to pyruvate and restores PEP via pyruvate, orthophosphate dikinase (PPDK) (Fig. 1).

**Figure 1.**
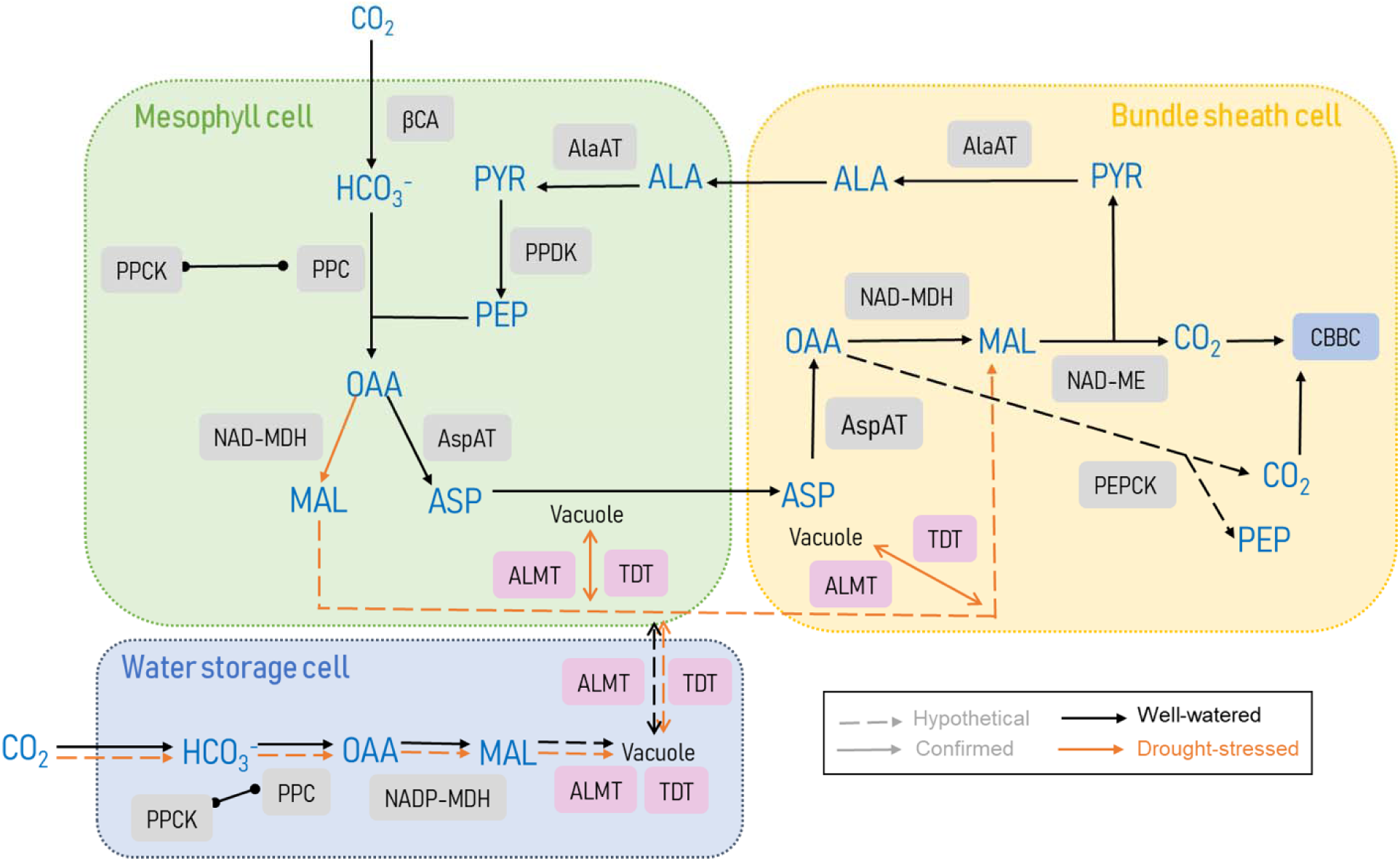
Schematic for the C_4_-CAM cycle in *P. oleracea* based on Ferrari & Reyna-Llorens (2026). Under well-watered conditions, the plant operates a dual-cell C_4_ cycle of subtype NAD-malic enzyme. Following drought stress, CAM is induced and since it involves movement of malate across different cell types, which is unsual for a CAM plants, this is defined as a dual-cell CAM mechanism, so far onl described in *Portulaca* species. Black lines indicate well-watered and orange, drought-stress. Continuous lines are steps confirmed empirically, whereas dashed lines are considered likely. Enzymes are indicated in grey boxes. Metabolites are written in blue. Abbreviations: βCA - β-carbonic anhydrase; PPC - phosphoenolpyruvate carboxylase; PPCK – PPC kinase; PEP –phosphoenolpyruvate; OAA – oxalaceate; MAL – malate; ASP – aspartate; NAD-MDH - NAD-malate dehydrogenase; NAD-ME – NAD malic enzyme; AspAT - aspartate aminotransferase; aluminum-activated malate transporter – ALMT; TDT - tonoplast malate/fumarate transporter; CBBC – Calvin-Benson-Bassham cycle; ALA – alanine; alaAT - alanine aminostransferase; PEPCK - phosphoenolpyruvate carboxykiane; PYR – pyruvate.

Once drought stress starts, CAM is gradually induced with specific upregulation of a CAM-specific PPC gene copy (*PPC-1E1c*), which is active during nighttime and stores CO_2_ as malate in the vacuoles of mesophyll cells (Christin *et al*., 2014, Ferrari *et al*., 2020a). Malate might also accumulate in water storage and bundle sheath cells (Ferrari & Reyna-Llorens 2026, Fig 1a). The C_4_ to CAM change also involves the change in peak transcript abundance of the enzyme responsible for activating PPC either during daytime (for C_4_) or nighttime (for CAM). This activation happens via phosphorylation by PPC kinase (PPCK) (Hartwell *et al*., 1996, 1999; Ferrari *et al*., 2020a). Overall, *P. oleracea* operates a dual-cell CAM mechanism, producing malate in one cell type (mesophyll or water storage cells) and shuttling it to bundle sheath cells during the day for decarboxylation, comprising an innovative photosynthetic system (Lara *et al*., 2004; Moreno-Villena *et al*., 2022).

Two aluminum-activated malate transporter (ALMT) gene copies have also been shown to be more expressed either under well-watered (*ALMT-12E.1*) or drought stress (*ALMT-12E.2*) conditions and have been used as markers for either C_4_ or CAM (Ferrari *et al*., 2020a,b). Although the exact identity of the malate transporter involved in export and import from the vacuole has not yet been identified, the involvement of ALMTs is taken into consideration due to previous differential gene expression evidence (Ferrari *et al*., 2020a). The tonoplast malate/fumarate transporter (TDT) is another candidate possibly involved in malate movement in and out of the vacuole.

Contrasting to strong CAM plants like cacti, CO_2_ assimilation at night only rarely occurs in *P. oleracea*, with malate formation relying on respiratory CO_2_ recycling and/or early morning bursts of CO_2_ uptake, both of which characterize a type of weak, facultative CAM referred to as CAM cycling (Koch & Kennedy 1980, Winter & Holtum 2014, Ferrari *et al*. 2020b). For this reason and especially in the case of weak CAM, gas exchange is not diagnostic of CAM (Ferrari et al., 2020b). Moreover, stable carbon isotope ratios (δ^13^C) have already been shown to vary drastically and provide ambiguous measurements even in strong CAM plants (Winter & Holtum 2002, Messerschmid et al., 2021). Hence, the best method to assess facultative or weak CAM is monitoring circadian changes in titratable acidity, which can be time consuming and lab-intensive, but is able to efficiently identify even slight variations in organic acid accumulation, indicating CAM induction (Winter & Smith, 2022).

In addition to titratable acidity and especially for C_4_-CAM in *Portulaca*, previous transcriptomic studies have already characterized comprehensively the global profile for core CCM genes during the 24h cycle (Ferrari et al. 2020a). This allowed the selection of specific key genes and time points for monitoring the switch between C_4_ and CAM. Furthermore, at least nine signalling genes including transcription factors (TFs) and hormone-related genes for both C_4_ and CAM have also been validated in *P. oleracea* (Ferrari et al., 2022). Overall, gene expression combined to titratable acidity serves as functional diagnostic parameters to monitor CCM changes in *Portulaca* (Ferrari *et al*., 2020a,b; Gilman *et al*., 2022; Moreno-Villena *et al*., 2022; Ferrari *et al*., 2022).

Facultative CAM is an extremely plastic physiological trait, and even though it is surely induced by drought in *P. oleracea*, there are several reports of other abiotic factors inducing it in C_3_ species (Freschi & Mercier, 2012). Salt stress is another well-studied stress factor for CAM induction, especially in the C_3_ species *Mesembryanthemum crystallinum*, commonly known as ice plant. While previous studies did not show CAM induction under salt stress alone in *P. oleracea* (Habibi, 2020), in *P. grandiflora* and *P. molokiniensis* it was confirmed (Bakpa *et al*., 2026). Overall, CAM responses according to nutrient availability seem to be group-specific and are still poorly understood (Pereira & Cushman, 2019).

Nitrogen (N) is essential for plant homeostasis as it is an integral constituent of protein and nucleic acids (de Bang *et al*., 2020). It is mostly taken up as NO_3_^−^ and NH_4_^+^ in inorganic forms (Ganeteg *et al*., 2017), depending upon soil structure, composition, pH, among other features (Marschner, 2012). Most available CAM studies focus on these two ions and although there can also be organic N sources, these are more relevant for epiphytic species and will not be discussed here. Among the few studies available, different plant lineages showed specific preferences or toxicity responses according to the N source. In the well-studied CAM group of *Kalanchoë*, species *K*. *blossfeldiana K. daigremontiana, K. laxiflora,* and *K. delagoensis* maintained and increased CAM activity when exposed to 1-10 mM NO_3_^−^, while even 1 mM NH_4_^+^ was toxic (Ota, 1988; Ota *et al*., 1988, 1991; Pereira *et al*., 2017). The epiphyte *Guzmania monostachia,* on the other hand, under drought stress showed increased CAM activity when grown with 5 mM NH_4_^+^ as opposed to NO_3_^−^ (Pereira *et al*., 2018), whereas N deficit even under well-watered conditions induced CAM (Rodrigues *et al*., 2014). Furthermore, facultative C_3_-CAM *Calandrinia polyandra* supplemented with 20 mM KNO_3_ repressed CAM activity (Winter & Holtum, 2011).

Phosphorus (P) deficiency is the second most limiting for plant growth, since P is part of essential biomolecules such as nucleic acids and ATP (Schachtman *et al*., 1998). Consequently, P deficiency causes a reduction in daytime CO_2_ assimilation and photosynthetic output (Carstensen *et al*., 2018). P is taken up as H_2_PO_4_ and HPO_4_^2-^ depending on the soil pH (de Bang *et al*., 2020). Under N+P deficiency, *M. crystallinum* showed higher CAM activity, which was also connected to a lower stomatal conductance (Paul & Cockburn, 1990). In *Clusia minor*, the only tree capable of performing CAM, N+P deficiency resulted in higher CAM activity under drought stress (Maiquetía *et al*., 2009). However, the isolated effects of P deficiency on CAM are not well understood yet.

CAM interactions with four other macronutrient elements are even less studied. First, potassium (K) is important for osmotic regulation and stomatal movements, with K^+^ found in vacuoles to provide turgor (de Bang *et al*., 2020). Although K can mitigate the effects of drought stress via sustaining assimilation rates in C_3_ crops (Cakmak, 2005), not much is known about its influence over CAM induction (Pereira & Cushman, 2019). Second, calcium (Ca) is overall important for cell walls and membranes, but cytosolic, ionic Ca is essential for transducing signals detected from the environment, having already been implied as a messenger also during facultative CAM induction in *M. crystallinum* (Taybi & Cushman, 1999). In well-watered *G. monostachia*, no CAM expression was induced solely under P, K or Ca deficiency (Rodrigues *et al*., 2014). Third, magnesium (Mg) is central in the chlorophyll molecule ring and shows strong electropositivity, affecting several enzyme catalytic reactions and showing a clear chlorotic phenotype in leaves during deficiency (de Bang *et al*., 2020). Lastly, sulphur (S) is usually taken up as inorganic sulphate (SO_4_^2-^) and its deficiency directly impacts leaf morphology (Marschner, 2012). Still, no studies have been done so far that specifically report CAM-associated changes during Mg and S deficiency.

Therefore, there is a lot of room for exploring the effects of individual nutrients over CAM expression, especially in facultative CAM plants. Moreover, as reviewed in Rodrigues *et al*. (2014) and Pereira & Cushman (2019), few studies assessing CAM modulation according to mineral nutrition monitored transcript abundance for key CCM genes as markers for CAM induction. For C_4_-CAM plants there is an even bigger knowledge gap as no studies have evaluated nutritional impacts in their metabolism (Pereira & Cushman, 2019), although good gene markers for C_4_ and CAM are available and have been validated (Ferrari *et al*., 2020a, 2020b, 2022).

Long term deficiencies have drastic effects over plant metabolism (de Bang *et al*., 2020), but short-term nutrient withdrawal as a treatment can easily provide insights into a fast-responding metabolic adaptation such as facultative CAM. In this context, the goal of this work was to investigate CAM-related physiological and transcriptional changes in *P. oleracea* studying the nutrients N, P, K, Mg, Ca, and S individually, but also in combination with drought stress. Salt stress was also included in the current work. Hydroponics was used to precisely control nutrient concentrations and PEG addition to the medium established lower osmotic potentials to mimic drought stress. Briefly, results showed that NO_3_^−^ deficiency and salt stress were potent CAM effectors, P deficiency showed a clear trend in inducing CAM, and Ca deficiency strikingly prevented CAM induction. Overall, these findings are extremely relevant in the context of dissecting the facultative C_4_-CAM coevolution and understanding the mechanism and regulation of a dual-cell CAM operation. This knowledge might also contribute to informing future endeavors of engineering C_4_ crops for facultative CAM.

## Materials and Methods

### Plant material and growth conditions

Seeds of *P. oleracea* were purchased from Kiepenkerl GmbH (Everswinkel, North Rhine-Westphalia, Germany). Seeds were germinated in a box with wet paper towel and a transparent lid for four days and transferred to 3 L pots filled with half strength Hoagland nutrient solution. Each pot initially contained 10 plants, and the solution was changed weekly. After two weeks, full strength Hoagland nutrient solution was used (1mM KH_2_PO_4_; 5mM KNO_3_; 5 mM Ca(NO_3_)_2_.4H_2_O; 2 mM MgSO_4_.7H_2_O; 0.2 mM FeEDTA; 1 µM H_3_BO_3_; 1 µM MnSO_4_. H2O; 0.2 µM CuSO_4_.5H_2_O; 0.01 µM (NH_4_)_6_Mo_7_O_24_.4H_2_O; 1 µM ZnSO_4_.7H_2_O) and the solution was changed 2x per week. The plants were kept in a walk-in climate chamber with a photoperiod of 12h light / dark and with temperatures of 28 / 22°C and relative humidity 60 /80 % at day / night (Fig. S1) The light intensity at plant level was at least 250 μmol m^−2^ s^−1^.

### Nutrient deficiency treatments

After one month growing, eight plants per group were transferred to 1 L pots and exposed to different nutritional treatments for one week, with solutions modified from Hoagland & Arnon (1950) (Table S1). Every treatment was applied with and without polyethylene glycol (PEG). The amount of PEG 8000 added per pot was calculated to establish an osmotic stress of −0.8 MPa according to Michel (1983). The following nutritional treatments were used: control (full strength Hoagland solution); PEG control (the same composition as the control but with PEG); salt treatment (200mM NaCl added to the control solution); NO_3_^−^deficient; with NH_4_^+^; half NO_3_^−^ and half NH_4_^+^; K deficient; Mg deficient; P deficient; S deficient; and Ca deficient (Table S1). The solution was changed two times during the treatment.

After one week, sampling of all treatments was performed at the same time: in the morning (07:00 h, one hour after the lights were on) and afternoon (17:00 h, one hour before the lights were off). Adult leaves were flash-frozen in liquid nitrogen and kept in a −80 °C freezer until use. Four biological replicates were collected alternately (e.g., first for all treatments, followed by the second, then the third, and lastly the fourth) across all treatments to avoid interference of the biological clock.

### Titratable acidity

Leaf samples were powdered without thawing and 200 mg were aliquoted into 1.5 mL tubes. Organic acids were extracted twice in 1.5 mL of 80% (v/v) methanol for 20 min at 80°C after 10min vortexing, and the supernatants were recovered twice by centrifugation (maximum speed, 5 min, 25°C), yielding 3 mL of extract per sample. Samples were titrated to pH 8 with small increments of 0.002 NaOH using phenolphthalein as an indicator. Diel acidity changes (ΔH^+^) were calculated subtracting morning (07:00 h) and afternoon (17:00 h) acidity values. A positive value indicated nocturnal accumulation of acids. Samples were normalized by fresh weight (FW).

### Osmotic potential (Ψs)

50 mg of ground plant tissue was used for collecting the sample sap after full speed centrifugation and read using the Osmomat 3000 Basic from Gonotec^©^. Osmolality values, provided as mOsmol/kg by the osmometer, were then converted MPa using the Van’t Hoff equation: Ψs = −CiRT, where C is the molar concentration of the solutes (molarity = moles L^−1^), i is the osmotic coefficient (considered 1 for molecules that do not dissociate in solution), R is the gas constant (8.31 J K^−1^ mol^−1^), and T is the absolute room temperature (K = 25°C + 273°).

### RNA extraction and quantitative real-time PCR

Total RNA was extracted from 80 mg of frozen powdered material using the ReliaPrep RNA Tissue Miniprep System (Promega) according to Ferrari *et al*. (2020c) in four biological replicates. RNA concentration was measured using a BioTek Epoch Microplate Spectrophotometer with the BioTek Take3 module and the Take3 Trio Microvolume Plate (Agilent, Santa Clara, United States). Purity was confirmed by maintaining absorbance ratios within 1.8 < A260/280 < 2.2 and 1.6 < A260/230 < 2.2. The iScript™ gDNA Clear cDNA Synthesis Kit (Bio-rad,Hercules, USA) was used for DNA digestion and cDNA synthesis. Quantitative real-time PCR (qPCR) was performed in 10 µL reactions containing 5 µL SYBR^®^ Green Supermixes for Real-Time PCR (Bio-rad, Bio-rad, Hercules, USA), 2 µL of diluted cDNA and 500 nM of each primer. All biological replicates were measured in multiple technical replicates for each gene analysed. The thermal cycling programme was set to: initial denaturation at 95 °C for 10 min; 40 cycles of 95 °C for 15 s, 60 °C for 30 s and 72 °C for 30 s; and a melting-curve analysis at the end of each run. Relative transcript abundance of the genes was calculated using the 2^−ΔΔCT^ method (Livak and Schmittgen, 2001) and the primer efficiency was calculated using LinRegPCR (Untergasser *et al*., 2021). All primers used are listed in Table S2, and gene names for core CCM components were followed as available in the literature (Christin *et al*., 2014, 2015, Ferrari *et al*., 2020a, b, 2022, Gilman *et al*., 2022; Moreno-Villena *et al*., 2022). Gene names for the signalling candidates were adopted considering their *Arabidopsis thaliana* annotation as previously published (Ferrari *et al*., 2022).

### Statistical testing

All statistical analyses and data plotting were performed using R (version 4.5.1; R Core Team, 2021) via RStudio (version 2025.05.1). Normality of the data was assessed using the Shapiro–Wilk test, and variance homogeneity was evaluated with Levene’s test. Appropriate tests were used in each case and according to the number of means being compared. In general, comparisons between two means were made using one of the following: two-sample t-test (for normally distributed, homoscedastic data); Welch’s t-test (for normally distributed, heteroscedastic data); or Wilcox test (for non-normal, homoscedastic data). Comparisons for more than three means were made using one of the following: one way ANOVA with Tukey’s test as posthoc test (normal, homocedastic); one way Welch ANOVA with Games-Howell as posthoc test (normal and heterocedastic); Kruskal-Wallis and Dunn’s test as posthoc test (non-normal and homocedastic). If the data were non-normal and heteroscedastic, transformations (square root or logarithmic) were applied and tests were repeated.

## Results

Plants were grown hydroponically first for two weeks using half-strength Hoagland’s solution, and later two more weeks using full-strength (Fig. S1). The plants were divided into 10 treatments without PEG: control (full-strength Hoagland solution); NO_3_^−^ deficient (−NO_3_^−^), with NH_4_^+^ (NH_4_^+^), half NO_3_- and half NH_4_^+^ (NO_3_^−^ + NH_4_^+^), Mg deficient (−Mg), P deficient (−P), Ca deficient (−Ca), K deficient –(K), S deficient (−S) and salt stress. Complementarily, a second group with the same 10 treatments received PEG to mimic drought stress responses by lowering the Ψs of the solution. This was given in an independent replicate for all treatments, and the control solution with PEG is referred to as just “PEG” throughout the text and images.

### Nutritional treatments significantly affected Ψs and ΔH^+^ with and without PEG

Ψs measurements in the treatments without PEG were not significantly different from control (−0.367 MPa ± 0.02), except for −S (−0.441 MPa ± 0.015) and salt stress (−0.697 MPa ± 0.013) (Fig. 2a, Table S3). When PEG was added, values were naturally more negative in most treatments, similarly to the PEG control (−0.736 MPa ± 0.03), except for −NO_3_^−^ (−0.518 MPa ± 0.015), −Mg (−0.597 MPa ± 0.010), and −P (−0.563 MPa ± 0.06) treatments, which showed significantly more positive Ψs values, and salt stress showed a significantly more negative value (−0.920 MPa ± 0.04) (Fig 2b, Table S3).

**Figure 2.**
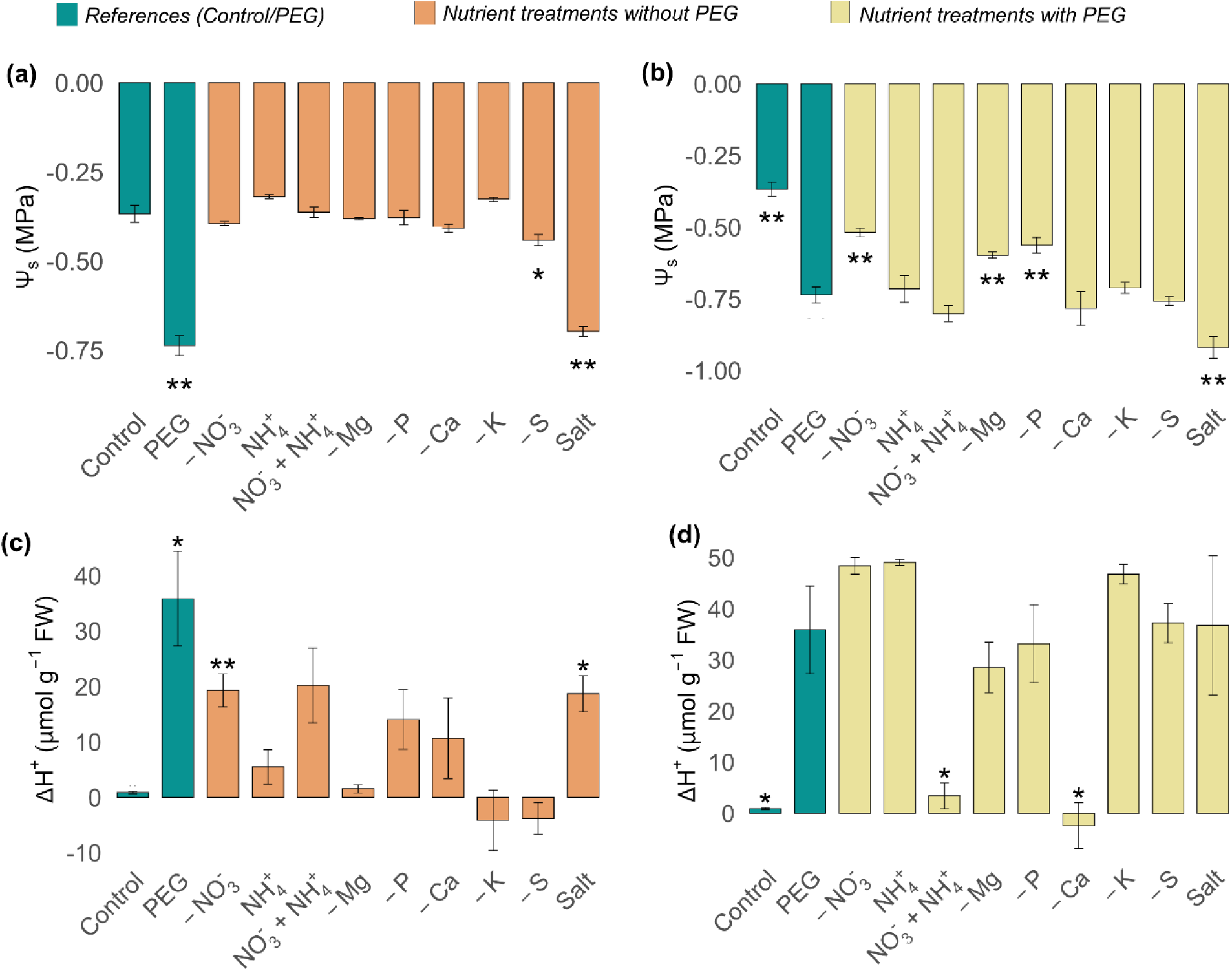
Physiological changes associated with the photosynthetic transition of C_4_ to CAM in *Portulaca oleracea.* Plants were grown hydroponically for a month in Hoagland’s solution and given 10 different nutritional treatments with and without PEG. (a-b) Osmotic potential (Ψs) in treatments without (a) and with (b) addition of PEG. (c-d) Titratable acidity difference when comparing morning (07:00 h) and afternoon (17:00 h) samples (ΔH^+^) in treatments without (a) and with (b) addition of PEG. Data are means and standard error of at least three biological replicates. Asterisks indicate pairwise statistical comparisons between treatments and either the references, which are the control without PEG in orange plots or the PEG control in yellow plots (*p<0.05; **p<0.01). Values used for plotting are given in Supplementary Tables S3 and S4.

Titratable acidity was monitored at dawn and dusk and the difference between those measurements representing the circadian variation in organic acid accumulation is referred here as ΔH^+^. ΔH^+^ was measured in all treatments with and without PEG (Fig. 2c-d, Table S4). The control group showed very low ΔH^+^ values (1.7 µmol H^+^ g^−1^ FW ± 0.18), (Fig 2c), while the PEG group reached 35.9 µmol H^+^ g^−1^ FW ± 7.4 one week after the start of the treatment. Without PEG, only −NO_3_^−^ (19.3 µmol H^+^ g^−1^ FW ± 2.57) and salt stress (18.7 µmol H^+^ g^−1^ FW ± 2.81) showed ΔH^+^ significantly higher than control, although the following treatments showed a positive trend: NO_3_^−^ + NH_4_^+^ (20.2 µmol H^+^ g^−1^ FW ± 5.85), −P (14.1 µmol H^+^ g^−1^ FW ± 4.65) and −Ca (10.7 µmol H^+^ g^−1^ FW ± 6.3) (Fig 2d, Table S2). With PEG, most nutrient treatments showed high positive ΔH^+^ values, except NO_3_^−^ + NH_4_^+^ (3.387 µmol H^+^ g^−1^ FW ± 2.22), and −Ca (−2.430 µmol H^+^ g^−1^ FW ± 3.9), which were significantly lower (Fig. 2d, Table S4).

### The transcription of C_4_, CAM, and signalling genes clearly revealed the impact of nutrition over CCMs

Next, relative expression was calculated for key genes involved in the core reactions and regulation of C_4_-CAM (Ferrari *et al*., 2020a, 2020b, 2022; Table S5). Figure 3a shows the assessed genes in the C_4_-CAM pathway of *P. oleracea*. βCA dissociates CO_2_ into HCO_3_^−^. β*CA-2E3* was expressed at similar levels in −NO_3_^−^, NH_4_^+^, and −P, while it was significantly downregulated in NO_3_^−^ + NH_4_^+^, −Mg, −Ca, −K, and −S. Treatments with added PEG were similarly downregulated for β*CA-2E3* relative expression, but specifically in −Ca, −K and −S levels were significantly lower than PEG (Fig. 3b).

**Figure 3.**
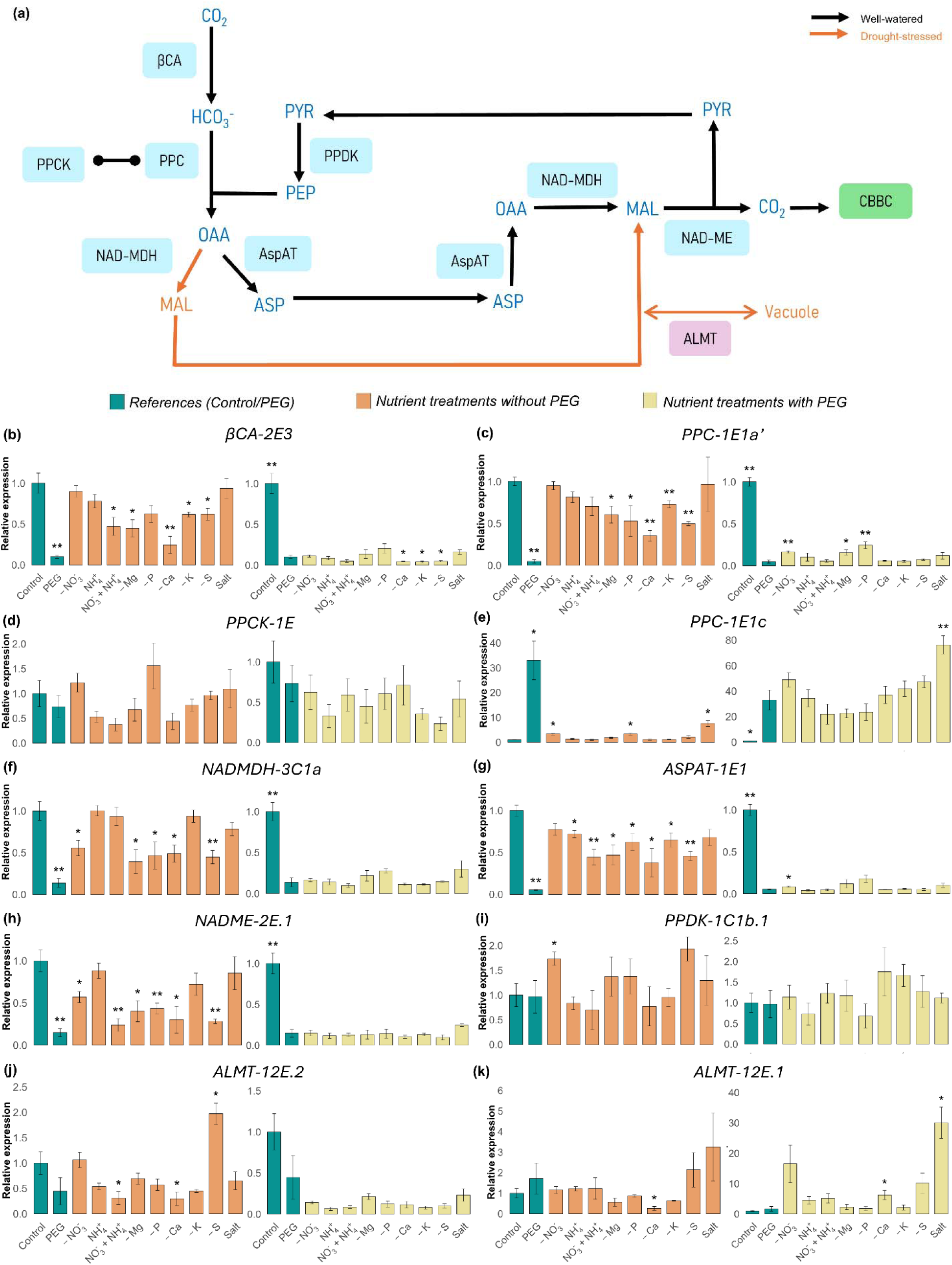
Relative expression changes in genes involved in core reactions of the C_4_-CAM pathway. (a) Schematic overview of the pathway considering well-watered and drought-stressed conditions. (b)-(k) Relative transcript abundance of β*CA-2E3* (b), *PPC-1E1a’* (c), *PPCK-1E* (d), *PPC-1E1c* (e), *NADMDH-3C1a* (f*), ASPAT-1E1* (g), *NADME-2E.1* (h), *PPDK-1C1b.1* (i), *ALMT-12E.2* (j), *ALMT-12E.1* (k). Data = mean and standard error of at least 3 biological replicates. Mean relative expression in all samples (with and without PEG) was normalized against the well-watered control samples without PEG in the morning for all genes except *PPC-1E1c’*, for which afternoon (17:00 h) was used. Asterisks indicate pairwise statistical comparisons between treatments and either the references, which are the control without PEG in orange plots or the PEG control in yellow plots (*p<0.05; **p<0.01). Abbreviations: βCA - β-carbonic anhydrase; PPC - phosphoenolpyruvate carboxylase; PPCK – PPC kinase; PEP –phosphoenolpyruvate; OAA – oxalaceate; MAL – malate; ASP – aspartate; NAD-MDH - NAD-malate dehydrogenase; NAD-ME – NAD malic enzyme; AspAT - aspartate aminotransferase; aluminum-activated malate transporter – ALMT; TDT - tonoplast malate/fumarate transporter; CBBC – Calvin-Benson-Bassham cycle; ALA – alanine; alaAT - alanine aminostransferase; PEPCK - phosphoenolpyruvate carboxykiane; PYR – pyruvate. Values used for plotting are given in Supplementary Table S5.

PPC combines PEP and HCO_3_^−^ to form oxalacetate. *PPC-1E1a’*, involved in C_4_, was downregulated in treatments −Mg, −P, −Ca, −K, −S without PEG, but sustained levels similar to the control in −NO_3_^−^, NH_4_^+^, NO_3_^−^ + NH_4_^+^, and salt stress, although for the latter with a high variation (Fig. 3c). With the addition of PEG, it was downregulated in all treatments, however −NO_3_^−^, −Mg and −P were slightly upregulated compared to the PEG control (Fig. 3c). *PPC-1E1c*, involved in CAM, was significantly upregulated in −NO_3_^−^, −P and salt stress without PEG, and with PEG transcripts were upregulated from 33- to 76-fold in all nutrient treatments compared to the control without PEG (Fig. 3e). PPCK (*PPCK-1E*) as the activator of PPC during the day in C_4_ and the night in CAM was not significantly modulated in any treatment, possibly since the sampled times (07:00 and 17:00) were not optimal for assessing differences in its expression profile (Fig. 3d).

NAD-MDH interconverts oxalacetate and malate, whereas NAD-ME decarboxylates malate to release CO_2_ (Fig. 3a). Here, similar patterns were observed for *NADMDH-3C1a* and *NADME-2E.1* across treatments (Fig. 3f,h). Without PEG, transcripts for both genes showed a significant downregulation in treatments −NO_3_^−^, −Mg, −P, −Ca, and −S. The only exception was NO_3_^−^ + NH_4_^+^, which was also downregulated for *NADME-2E.1* without PEG (Fig. 3h). With PEG, all treatments showed a significant and stronger downregulation (Fig. 3f,h). AspAT is responsible for interconverting between oxalacetate and aspartate. *ASPAT-1E1* was significantly downregulated in all treatments except −NO_3_^−^ and salt stress without PEG, and in −NO_3_^−^ with the addition of PEG (Fig. 3g). PPDK is responsible for regenerating PEP from pyruvate (Fig. 3a). An upregulation in relative expression for *PPDK-1C1b.1* was observed only in −NO_3_^−^ without PEG, all other treatments were not significantly different than the respective controls (Fig. 3i).

The transporter ALMT is putatively involved in malate movements in and/or out of the vacuole (Fig. 3a). *ALMT-12E.2* is expressed under well-watered conditions and was present at similar levels in all treatments without PEG, except in NO_3_^−^ + NH_4_^+^ and −S, while it was downregulated in all treatments with PEG (Fig. 3j). On the other hand, *ALMT-12E.1*, which is potentially involved in CAM, was not modulated with the addition of PEG in most treatments, except in −Ca (also without PEG), and salt stress (Fig. 3k).

The last step in the current analysis involved monitoring the expression of genes previously confirmed to be a part of the regulation and signalling of CAM in C_4_ *P. oleracea* (Ferrari *et al*., 2022). Nuclear Factor Y (NFY) is a family of transcription factors commonly involved in drought stress responses (Amin *et al*., 2019), and *NFY SUBUNIT C4* (*NFYC4*) and *NFY SUBUNIT A7*. Here, *NFYC4* was in general downregulated in all treatments with the addition of PEG as opposed to without it (Fig. 4a). *NFYA7*, on the other hand, was upregulated over 70-fold in all PEG treatments compared to the control without PEG (52-fold in −Ca). However, even without PEG, it was already upregulated 9.5-fold in −NO_3_^−^, 4.38-fold in NO_3_^−^ + NH_4_^+^, and 4.36-fold in −P (Fig. 4b). *TRANSPARENT TESTA 8 (TT8)* was upregulated in a similar manner in all treatments with PEG, but significantly upregulated in NH_4_^+^, NO_3_^−^ + NH_4_^+^, and −K (Fig. 4c). *TT8* was not modulated in treatments with PEG, although it was more expressed than compared to treatments without PEG (Fig. 4c). Cis-epoxycarotenoid dioxygenase (NCED), specifically *NCED3*, is responsible for ABA synthesis (Okamoto *et al*., 2011). Without PEG, *NCED3* was upregulated ca. 4-fold in treatments −Mg, −P and −K, and also in NH_4_^+^, −K and salt stress with PEG (Fig. 4d). *ABA RESPONSIVE ELEMENT-BINDING FACTOR 2* (*ABF2*) was upregulated in treatments −Mg, −Ca, −K, −S and salt stress without PEG, and −K and salt stress with PEG (Fig. 4e). Lastly, transcription factor *HOMEOBOX* 7 (*HB7*) is highly responsive to drought stress (Valdés *et al.,* 2012) and was suggested as a CAM marker in *P. oleracea* (Ferrari *et al*., 2022). Here, *HB7* was upregulated thousands of times in all treatments with PEG, but also without PEG 71-fold in −NO_3_^−^, 9-fold in NH_4_^+^, 67-fold in −Mg, 24-fold in −P, and 51-fold in −Ca (Fig. 4f).

**Figure 4.**
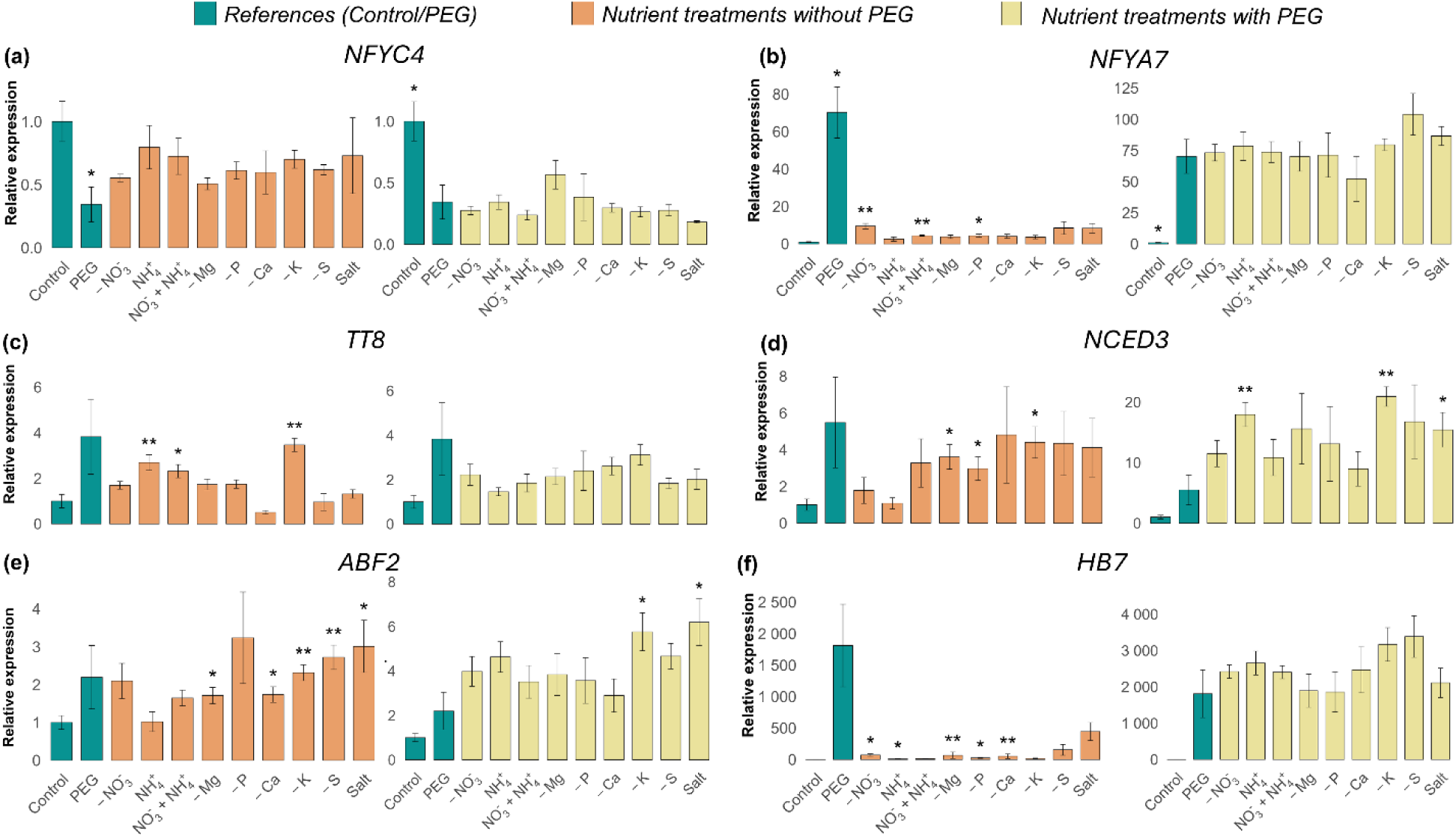
Relative expression of genes involved in the regulation and signaling of the CAM switch in C_4_ *Portulaca oleracea*. (a) *NUCLEAR FACTOR Y SUBUNIT C4* (*NFYC4*), (b) *NUCLEAR FACTOR Y SUBUNIT A7* (*NFYA7*), (c) *TRANSPARENT TESTA 8 (TT8), (d)* CIS-EPOXYCAROTENOID DIOXYGENASES (NCED3), (e) *ABA RESPONSIVE ELEMENT-BINDING FACTOR 2* (*ABF2*)*, (f) HOMEOBOX* 7 (*HB7*). Mean relative expression in all samples (with and without PEG) was normalized against the well-watered control samples without PEG in the afternoon (17:00 h) for all genes except NFYC4, for which morning (07:00 h) was used. Asterisks indicate pairwise statistical comparisons between treatments and either the references, which are the control without PEG in orange plots or the PEG control in yellow plots (*p<0.05; **p<0.01). Values used for plotting are given in Supplementary Table S5.

## Discussion

Photosynthesis as the process of generating energetic molecules (NADPH and ATP) and photoassimilates is directly and indirectly affected by all macronutrient deficiencies in the short and long-term (Zenda *et al.,* 2021, Wang *et al*., 2019, Lu *et al*., 2025). Whereas long-term deficiency causes several known symptoms connected with the functional roles of each nutrient, short-term deficiencies have impacts in the molecular environment of the plant (de Bang *et al.,* 2020). Deficiency syndromes are well-studied in crops, which are mostly performing C_3_ but also frequently C_4_, reinforcing that it is timely to deepen our understanding of nutrition over less studied CAM plants (Pereira & Cushman, 2019). Although the attribute of inducing or not CAM can be considered a discrete trait, a continuum of CAM physiotypes exist across the tree of life (Edwards, 2023), complexifying the characterization of common responses to a nutrient deficiency.

In the present work, the C_4_-CAM plant *P. oleracea* was studied for CAM-related physiological and transcriptional responses according to individual, short-term nutrient deficiencies. From the measured parameters, changes in ΔH^+^ together with changes in transcript abundance for core CCM genes and transcription factors (TFs) were considered diagnostic criteria for a significant modulation in either C_4_ or CAM (Table 1). Overall, positive ΔH^+^ in nutritional treatments without PEG accompanied by upregulation of *PPC-1E1c* were considered diagnostic of CAM induction. At the same time, negative ΔH^+^ was considered absence of CAM activity and unchanged gene expression of C_4_-related genes indicated sustained C_4_ metabolism (Table 1). The controls with and without PEG showed clear and expected responses, with no CAM induction without PEG and clear CAM induction with PEG (Figs. 2-4, Table 1).

**Table 1.**
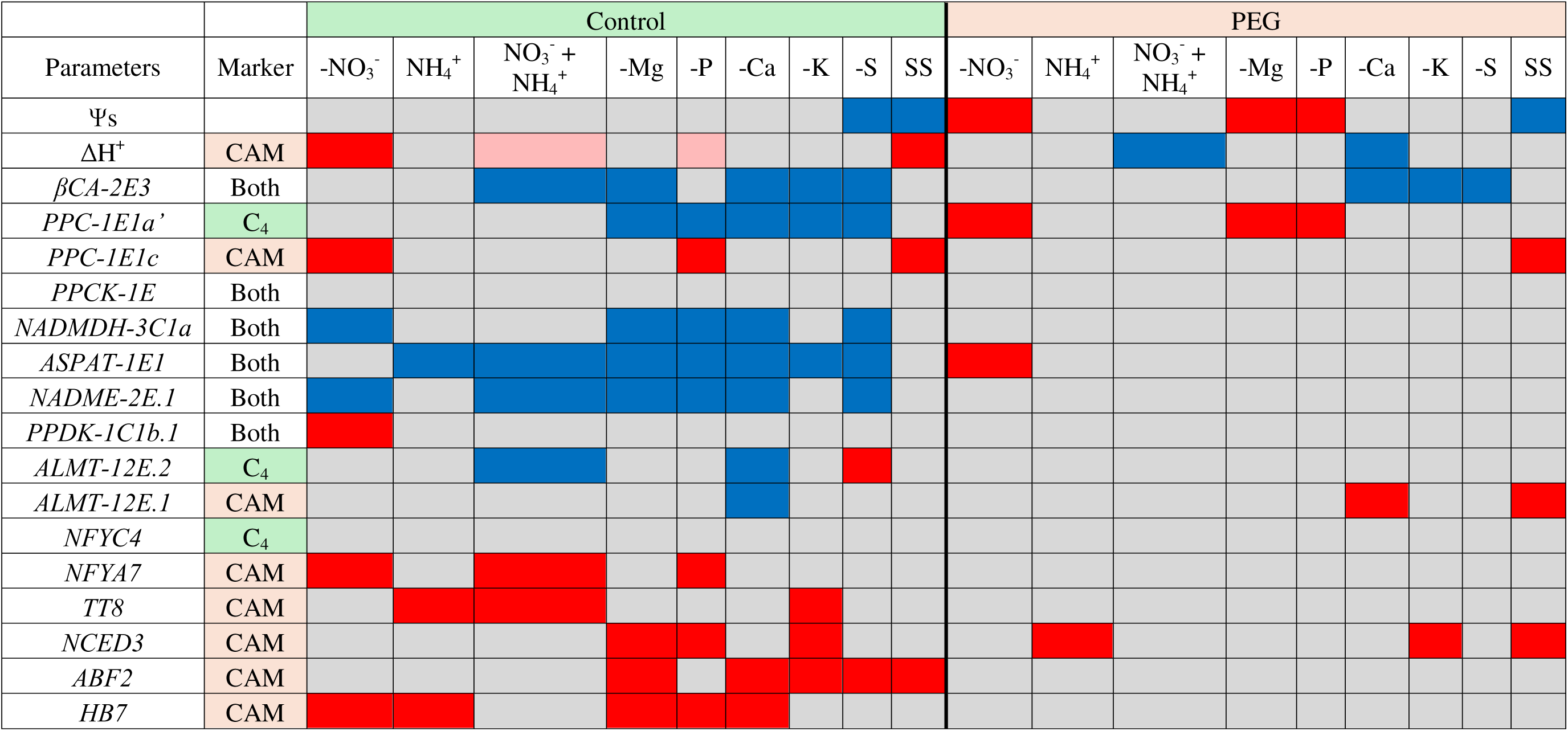
Visual summary of physiological and transcript relative abundance parameters monitored in leaves of *Portulaca oleracea* subjected to 10 different nutritional treatments with and without PEG. Abbreviations are defined in the text. Gray = non-significant values when compared to control or PEG. Red = significant upregulation. Blue= significant downregulation. Values used in this table are given in Supplementary Tables 3-5.

### Nitrate deficiency induces CAM in P. oleracea possibly via an ABA-independent route

The present work shows for the first time in a C_4_-CAM plant that nitrate deficiency (−NO_3_^−^) induced CAM even without the addition of PEG to the solution, since ΔH^+^ was significantly increased, there was higher relative expression of core CAM genes *PPC-1E1c* and *PPDK-1C1b.1*, and of CAM signaling TFs *NFYC7* and *HB7* (Table 1). Although *NADMDH-3C1a* and *NADME-2E.1* were downregulated in the −NO_3_^−^ treatment without PEG, *PPC-1E1a’* and *ASPAT-1E1* were still expressed in levels comparable to the C_4_ control. Since these two genes directly contribute to the flux of metabolites storing CO_2_ in C_4_-CAM, it might indicate that C_4_ and CAM were probably active at the same time. In addition, in −NO_3_^−^ with PEG, less negative values of Ψs and a significant increase in the relative expression of *PPC-1E1a’* were observed, suggesting a small rescue of C_4_ compared to the PEG control (Figs. 2-3; Table 1). Overall, this crosstalk between C and N metabolism represents a promising set up for further exploring and understanding C_4_ and CAM concomitant activation and compatibility in *Portulaca*.

Compared to the few other CAM groups that have been studied, it seems that a species-specific relationship between C and N sources exists. In constitutive CAM species from *Kalanchoë* and Cactaceae, supplementation with NO_3_^−^ resulted in increased CAM activity via higher acid production and PPC activity, which makes sense since these plants acquire most of their C via CAM (Winter *et al*., 1982, Nobel *et al*., 1983, Hartwell *et al*., 2016). However, in facultative CAM species such as *C. polyandra* and *G. monostachia*, increased NO_3_^−^ availability does not result in CAM induction (Winter & Holtum 2011, Pereira *et al*., 2018). Nevertheless, −NO_3_^−^ treatment with and without drought stress was shown to induce CAM in *G. monostachia* (Rodrigues *et al*., 2014), agreeing to the effects observed here for *P. oleracea*. In general plant metabolism, malate and NO_3_^−^ metabolism intertwine during daytime, when malate is produced to neutralize hydroxyl groups deriving from NO_3_^−^ assimilation, but this is not connected with CAM behaviour (Winter & Smith, 2022).

Here, for *P. oleracea*, a positive CAM response in −NO_3_^−^ plants without PEG was accompanied by a lack of modulation of *NCED3* and *ABF2*. These genes are involved in ABA biosynthesis and signaling, respectively, suggesting CAM induction without the mediation of ABA, although levels for this hormone were not quantified here. ABA has been shown successively to be a CAM effector hormone intrinsically connected to drought stress, but an ABA-independent route for CAM induction has been also suggested (Freschi & Mercier, 2012). Moreover, cytokinin is usually suggested as a CAM inhibitor (Pereira *et al*., 2013). Interestingly, tomato (*Lycopersicon esculentum*) starved from NO_3_^−^ for few hours showed a reduction in cytokinin concentration both in zeatin or zeatin riboside forms, which reduced leaf growth (Rahayu *et al*., 2005). Moreover, exogenous application of 6-Benzylaminopurine (BA, a synthetic cytokinin) or zeatin was shown to significantly reduce transcript abundance of the CAM-PPC (*PPC-1E1c*) in *P. oleracea* (Ferrari *et al*., 2022). Therefore, this could suggest that CAM induction in the −NO_3_^−^ treatment was more a result of reduced cytokinin signalling instead of direct ABA mediation, but further experiments quantifying these hormones are required.

### Ammonium is not toxic for P. oleracea and does not induce CAM

In the present work, replacing NO_3_^−^ by NH_4_^+^ as N source did not result in a toxicity response, since genes involved in the C_4_ metabolism remained unchanged (Fig. 3), NH_4_^+^ treatment with PEG enabled CAM induction comparable to the PEG control (Fig. 2), and there were no visual signs of necrosis on roots and leaves. At the same time, NH_4_^+^ did not induce CAM since there was no modulation of ΔH^+^ and *PPC-1E1c* mRNA abundance (Figs. 2-3). NH_4_^+^ toxicity symptoms may include stimulation of photorespiration (Britto & Kronzucker, 2002), which although not assessed here, was not enough to induce CAM in *P. oleracea*. Although C_4_ gene expression was not altered in the present work, plant height, leaves, and plant fresh biomass were reported to decrease as *P. oleracea* plants were exposed to a NH_4_^+^/total N ratio higher than 0.05 for ca. 20 days (Chrysargyris *et al*., 2023). This suggests a limit for NH_4_^+^ tolerance in the species.

The preference for different N sources appears to be species-specific in terms of CAM expression, since NH_4_^+^ is toxic for several *Kalanchoë* species but enhances CAM in the bromeliad *G. monostachia* (Ota, 1988; Ota *et al*., 1988, 1971; Pereira *et al*., 2017, 2018). In plant metabolism, whereas NO_3_^−^ needs to be converted to NH_4_^+^ to be metabolized and form amino acids, NH_4_^+^ is readily available. Among the reasons for NH_4_^+^ toxicity are a disturbance of the cell pH and a high energetic cost (1 NH_4_^+^ : 1 ATP) to pump ammonium out of the cells, which could also affect CAM responses (Pereira & Cushman, 2019).

In the present study, a trend to increased but non-significant ΔH^+^ when compared to the control was observed in the treatment with half nitrate and half ammonium (NO_3_^−^ + NH_4_^+^) without PEG (ΔH^+^ = 20.2 µmol H^+^ g^−1^ FW ± 7.46, Fig. 3). Even though it was not accompanied by significantly increased *PPC-1E1c* relative transcript abundance (Fig. 3, Table 1), signaling TFs *TT8* and *NFYC7* were significantly upregulated (Fig. 4). The opposite effect was observed in NO_3_^−^ + NH_4_^+^ with PEG, since ΔH^+^ was significantly reduced compared to the PEG control. Overall, these effects were not clear and, without PEG, they might have been caused merely by a reduction in NO_3_^−^ availability. Generally, for the commercial growth of *P. oleracea* in which big and healthy leaves are desired, optimal proportions of NO_3_^−^ and NH_4_^+^ were measured to be circa 3-14 mM and 0.4-0.8 mM, respectively (Chrysargyris & Tzortzakis, 2025).

### CAM induction during phosphate deficiency might alleviate high C_4_ demands for Pi

The phosphate deficiency (−P) treatment without PEG (Fig. 2) also showed a trend to increased ΔH^+^ 7 and, in this case, it was accompanied by significant increases in relative expression of *PPC-1E1c*, *NFYC7*, *NCED3*, and *HB7* (Figs. 3-4; Table 1). Without PEG, there was further downregulation of C_4_ genes *NADMDH-3C1a*, *ASPAT-1E1*, and *NADME-2E.1*, likely indicating a downregulation of C_4_ metabolism as CAM was induced. Indeed, *Flaveria* species as a C_4_ model were shown to experience stronger inhibition of photosynthesis under −P compared to C_3_ *Panicum* species assessed by gas exchange monitoring (Krone *et al.,* 2025).

Here, the −P treatment with PEG also resulted in a significant increase in Ψs and *PPC-1E1a’* relative mRNA abundance compared to the PEG control, similar to −NO_3_^−^ and −Mg (Figs. 2-3; Table 1). For *M. crystallinum*, pinitol was shown to accumulate in plants with induced CAM and under −P and −N, with pinitol potentially interfering with the osmotic adjustment of the chloroplast (Paul & Cockburn, 1989, 1990). The more positive values of Ψs in *P. oleracea* might have been the result of similar metabolites accumulating exclusively under PEG treatment and possibly making the higher C_4_-*PPC* relative transcript levels a consequence of a lower level of osmotic stress in the cell.

Limited previous evidence is available about CAM induction according to P availability, but the data indicate positive CAM activity during −P also showed an influence of −N for *M. crystallinum* (Paul & Cockburn, 1990) and of mycorrhization for *C. minor* (Maiquetía *et al*., 2009). As such, the present work appears to be one of the first to provide evidences of the effect of P nutrition individually on CAM. Inorganic phosphate (Pi) is needed for photophosphorylation in the Calvin-Benson-Bassham cycle and for the regulation of several enzymes part of CCM metabolism such as PPC by PPCK, making the daytime demand for P high (Rychter & Rao, 2005, Hartwell *et al*., 1996). Therefore, inducing CAM under −P and assimilating CO_2_ at night could reduce the competition for Pi during the day. In CAM, flux balance analysis highlighted the importance of Pi for the mitochondrial phosphate carrier (PiC, Pi/H^+^ symport) in allowing mitochondria to generate necessary energetic demands for the CCM functioning (Daems *et al*., 2024). Indeed, in C_4_ plants there is a high P demand, which has been considered one of the possible factors causing C_4_ to be less phenotypically plastic compared to C_3_ (Sage & McKown, 2006, Krone *et al*., 2025). However, if a C_4_ plant is able to induce CAM under −P and optimize Pi demands, this would certainly contribute to adapting more easily to unfavorable environmental conditions, which seems to be the case of *P. oleracea*.

### Calcium deficiency prevented CAM induction even in P. oleracea plants treated with PEG

Calcium deficiency (−Ca) significantly altered ΔH^+^ with and without the addition of PEG in *P. oleracea* (Table 1). Without PEG, higher ΔH^+^ was accompanied by downregulation of several C*_4_*-related genes and upregulation of *ABF2* and *HB7* TFs (Figs. 2-4, Table 1). More strikingly, PEG treatment showed a significant decrease in ΔH^+^ compared to PEG control (−2.4 µmol H^+^ g^−1^ FW ± 3.9), although without associated transcription pattern changes for core CCM genes (Table 1). Changes in levels of intracellular free Ca^2+^ are a potent signaling cue for multiple pathways at the same time, such as photosynthesis (Brand & Becker, 1984), phytohormone signaling, and stomatal movements (Weinl *et al*., 2008). In the Calvin-Benson-Bassham cycle, sedoheptulose-1,7-bisphosphatase (SBPase) and fructose-l,6-bisphosphatase (FBPase) are key enzymes regulated by Ca (Raines, 2005). Moreover, CCMs have multiple steps regulated via phosphorylation and dephosphorylation processes, which are commonly performed by Ca^2+^- and calmodulin-dependent protein kinases and phosphatases 1 and 2A. Interestingly, PPCK is a Ca^2+^ independent kinase so PPC would not be affected by −Ca directly (Hartwell *et al*., 1996). However, there seems to be an upstream step in signaling to PPCK that involves Ca^2+^, which is still needed for PPC activation as shown in C_4_ species *Digitaria sanguinalis* (Duff *et al*., 1995). This evidence would justify a downregulation in overall C_4_ metabolism under −Ca even without PEG as observed here.

Ca was shown to play a role in CAM signaling both via an ABA-dependent and independent routes (Freschi & Mercier, 2012). Treating *M. crystallinum* plants with ethyleneglycol-bis(aminoethyl ether)-N,N′-tetraacetic acid (EGTA), a calcium chelator, prevented accumulation of transcripts for *PPC1*, *GAPC1* and *MDH1* during salt stress (Taybi & Cushman, 1999). Furthermore, Taybi & Cushman (1999) also treated unstressed leaves with ionomycin, a Ca^2+^ ionophore, and observed an increase in *PCC* transcripts even without stress. In pineapple, *Ananas comosus*, an obligate CAM plant, the same treatment with ionomycin was able to induce PPC, MDH and PEPCK activities, whereas EGTA prevented it (Freschi *et al*., 2010). Very limited information is available about the ABA-independent route of CAM signaling, but it also relies on Ca^2+^ signaling as a trigger for *PPC* expression (Taybi *et al*., 2002). Thus, it seems plausible that −Ca with PEG would have prevented CAM induction in *P. oleracea* and this probably occurred in an ABA-independent way.

Lastly, other treatments −Mg, −K, and −S exhibited unclear responses that could not be directly related to changes in CAM but were probably a result of the plant’s suboptimal state after exposure to a macronutrient deficiency, leading to a downregulation in C_4_ genes in treatments without PEG (de Bang *et al.,* 2020, Table 1). In addition, a consistent and significant reduction in relative expression of β*CA-2E3* was observed in −K, −S and also −Ca treatments with and without PEG.

As an extra treatment in the current work, salt stress was included since it is a potent CAM effector in *M. crystallinum* (Winter & von Willert 1972). In *Portulaca*, a recent study confirmed CAM induction upon salt stress in pot-grown *P. grandiflora* and *P. molokiensis* (Bapka *et al*., 2026). The present work adds to this report with confirmation of strong and clear CAM induction in *P. oleracea* grown in hydroponics with and without PEG. We observed significant and positive ΔH^+^, *PPC-1E1c* transcript accumulation, and here there was probably signaling and involvement of ABA, since *NCED3* and *ABF2* were also induced (Table 1).

## Conclusion

*Portulaca* species operate the uncommon feature of alternating between C_4_ photosynthesis during optimal environmental conditions and CAM during drought stress, with *P. oleracea* being the most studied species in the group. This work provided the first overview of C_4_-CAM physiological and transcriptional responses during macronutrient deficiencies, an abiotic stress topic that is currently still poorly understood in the literature for C_4_ or CAM plants, but even more in C_4_-CAM plants. In *P. oleracea*, NO_3_^−^ deficiency was a potent CAM inducer even under optimal watering, which was probably not mediated by ABA and instead connected to a downregulation of cytokinin metabolism. Since C_4_ was still active in −NO_3_^−^, this opens a new promising opportunity for understanding the compatibility and coexistence of C_4_ and CAM independent of drought stress. Moreover, C_4_ plants have a high demand for P and the findings of the present study indicate that its availability likely plays a greater role in CAM activity than currently discussed. In addition, Ca deficiency inhibited CAM activity even during PEG treatment, reinforcing the importance of Ca^2+^ signaling in CCM metabolism, a currently understudied topic. Regarding specific responses, *P. oleracea* was able to induce CAM during salt stress alone and tolerated NH_4_^+^ as a N source without toxicity responses. Overall, these findings add another dimension of abiotic factors that can affect CCMs and offer a different perspective for studies of C_4_-CAM functioning and evolution. In addition, biosynthetic biology approaches seeking to engineer C_4_-CAM can benefit from a deeper understanding of how mineral nutrients affect the molecular modulation of this dual CCM in *P. oleracea* (Ferrari & Reyna-Llorens, 2026).

## Acknowledgements

I would like to thank the support of Prof. Klaus Dittert, Manavi Shrestha for temporally taking care of the plants, Ulf Jäger for the technical support, Tino Kreszies for helpful discussions, and Vinicius Daguano Gastaldi for revising the manuscript.

## Author contribution

RCF designed the project, acquired funding, did experimental data collection and analysis, and wrote the manuscript.

## Data availability

All data discussed here is available in supplementary material.

## Supporting Information

### Supplementary Tables

S1 Nutrient solutions modified from the original Hoagland’s solution.

S2 Primers used for relative gene expression analysis via qPCR of *P. oleracea* samples under different nutritional treatments with and without PEG

S3 Osmotic potential (Ψs) measurements for *P. oleracea* samples under different nutritional treatments with and without PEG

S4 Titratable acidity differences between morning (7:00) and afternoon (17:00) (ΔH^+^) for *P. oleracea* samples under different nutritional treatments with and without PEG

S5 Relative expression calculated using qPCR data for key genes involved in the core reactions and regulation of C_4_-CAM in *P. oleracea* samples under different nutritional treatments with and without PEG

### Supplementary Figure

Supplementary Figure S1. General overview of the experiment including 10 different treatments for nutrient availability in *Portulaca oleracea*. (a) Aspects of the plants before the start of the nutrient +/− PEG. (b) Detail of one pot.

