## Supplementary Figure 1 for "Differential modulation of crassulacean acid metabolism according to macronutrient deficiencies in C_4_ *Portulaca oleracea*"

Author: Renata Callegari Ferrari

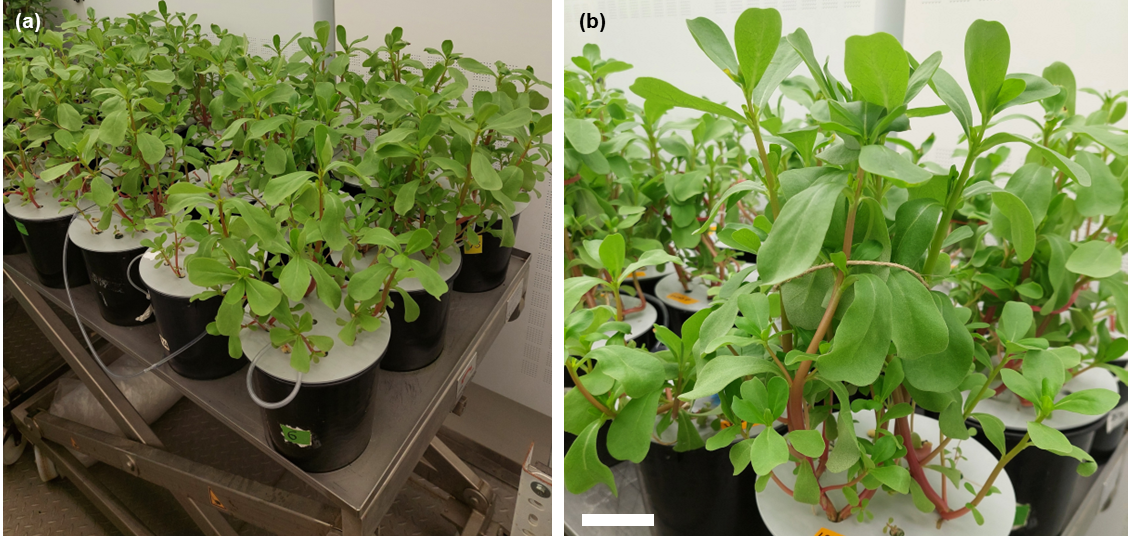

Supplementary Figure S1. General overview of the experiment including 10 different treatments for nutrient availability in *Portulaca oleracea*. (a) Aspects of the plants before the start of the nutrient +/- PEG. (b) Detail of one pot.
